# Regulated distribution of mitochondria in touch receptor neurons of *C. elegans* influences touch response

**DOI:** 10.1101/2020.07.26.221523

**Authors:** Anjali Awasthi, Souvik Modi, Sneha Hegde, Anusheela Chatterjee, Sudip Mondal, Eva Romero, Guruprasad R. Sure, Sandhya P. Koushika

**Author notes:** These authors contributed equally (Anjali Awasthi, Souvik Modi).

## Abstract

Density of mitochondria and their localization at specific sub-cellular regions of the neurons is regulated by molecular motors, their adaptors and the cytoskeleton. However, the regulation of the mitochondrial density, the positioning of mitochondria along the neuronal process and the role of axonal mitochondria in neuronal function remain poorly understood. This study shows that the density of mitochondria in *C. elegans* touch receptor neuron processes remains constant through development. Simulations show that mitochondrial positioning along parts of the neuronal process that are devoid of synapses is regulated. Additionally, we also demonstrate that axonal mitochondria are necessary for maintaining touch responsiveness.

## Introduction

Neurons are highly polarized cells containing multiple processes with high energy demands (Rossi and Pekkurnaz, 2019). They rely on the mitochondria not only to generate energy necessary for metabolism but also to buffer calcium (Duchen, 2012) and modulate reactive oxygen species turnover in neurons (Lopez-Fabuel et al., 2016). Thus, the distribution and maintenance of mitochondria throughout the neuron are important for neuronal function.

Neurons possesses long axons that can extend over a meter from the cell body. Cell body is the hub for majority of mitochondrial biogenesis and degradation, while after axon specification, mitochondria are transported into the axonal process (Chang et al., 2006; Saxton and Hollenbeck, 2012). Bi-directional transport mitochondria is correlated with axonal outgrowth and the mitochondrial motility is maintained throughout the axonal development (Morris and Hollenbeck, 1993; Ruthel and Hollenbeck, 2003). Mitochondria are found to be predominantly stationary in the axon and tends to present or accumulate, enriched/present in energy demanding areas (Chen et al., 2016). These regions include growth cones (Morris and Hollenbeck, 1993; Smith and Gallo, 2018), synaptic boutons (Levy et al., 2003), nodes of Ranvier (Zhang et al., 2010), axonal branches and dendritic shafts (Li et al., 2004; Ruthel and Hollenbeck, 2003). Mitochondrial positioning is thought to occur through a two-step process that involves transport using molecular motors (Schwarz, 2013) and local immobilization. Immobilization can be achieved either by detaching from cytoskeletal tracks or stalling by molecular anchors present in both cytoplasm and on mitochondrial surfaces (Kraft and Lackner, 2018). Mitochondrial positioning can also be regulated in an activity dependent manner, both in cultured neurons and *in vivo* (Stephen et al., 2015; Zhang et al., 2010). Further, studies from myelinated CNS fibers from brain slices of rodents, show that axonal mitochondria can pause or slow down preferentially in node–paranodal regions (Ohno et al., 2011). Demyelination is known to affect mitochondrial mobility as well as size of stationary mitochondria (Kiryu-Seo et al., 2010). Mitochondrial densities increase in dysmyelinated optic nerve axons and lesions of human brain sections (Hogan et al., 2009; Mahad et al., 2009). Similarly, slices obtained from patients suffering from multiple sclerosis showed chronically demyelinated spinal cord axons as well as a significantly reduced mitochondrial density (Dutta et al., 2006). These results suggest that mitochondrial motility and localization is regulated in both myelinated and non-myelinated neurons.

Studies from *C. elegans* sensory and motor neurons which lack a dedicated myelin sheath (White et al., 1986) indicate a constant density of mitochondria in the axonal processes, which is dependent on the mitochondrial trafficking machinery (Han et al., 2016; Morsci et al., 2016; Sure et al., 2018). However, it is still not clear how mitochondrial number and densities vary in neurons during development from larval stages to young adults. It is also unclear how mitochondria are distributed along the long neuronal process *in vivo*, essentially which are devoid of reported synapses or nodes of Ranvier. Mitochondrial distribution along the axonal process is critical for various cellular functions including energy production (Harris et al., 2012), calcium homeostasis (Billups and Forsythe, 2002), apoptosis (Frezza et al., 2006), synaptic transmission (Sun et al., 2013). Thus, it is essential to understand whether this distribution is developmentally regulated and whether alteration of the distribution can influence the animal behavior.

Using *C. elegans Touch* Receptor Neurons (TRNs) as a model system, we show that mitochondrial density in the axonal process remains constant throughout its larval development. Mitochondria are added proportionate to axon growth from 1st larval stage (L1) to young adult (1d adult), thus resulting in constant mitochondrial density. We also found that inter-mitochondrial distances show significant deviation from a uniform or random distribution. This non-random distribution is attained at 2^nd^ larval stage and remains unchanged up to adult stages. Our data also suggests that the distribution is independent of molecular motors and adapter proteins. Finally, we show that the presence of mitochondria along the neuronal process is important for gentle touch response. Animals with mutations in genes that affect mitochondrial density and distribution respond poorly to gentle touch stimulus. These results indicate that mitochondrial density *in vivo* regulated by cellular trafficking machinery and positioning of mitochondria along the neuronal processes is important for animal behavior.

## Results

### Mitochondrial density and distribution in TRN is regulated

*C. elegans* have six touch receptor neurons (Fig. 1*A*) and mitochondria are known to be present throughout the touch neuronal process (Fig. 1*B*). The density of mitochondria in a 1d adult is seen to be ~5 mitochondria per 100 μm in Anterior Lateral Microtubule (ALM) neurons (Morsci et al., 2016; Sure et al., 2018). We found that the density of mitochondria in adult Posterior Lateral Microtubule (PLM) (5.19±0.52/100 μm, Mean±SD) neurons, is similar to ALM (5.43±0.78/100 μm, Mean±SD) (Fig. 1*C*). This density remains constant throughout the development of *C. elegans* TRNs from the L1 larval stages to adult stages (Supplementary Fig. S1*A-C*). Additionally, significantly greater variance in the mitochondrial density is observed in L1 larval stage compared to other developmental stages (Supplementary Fig. S1*D*, Table 1).

**Figure 1.**
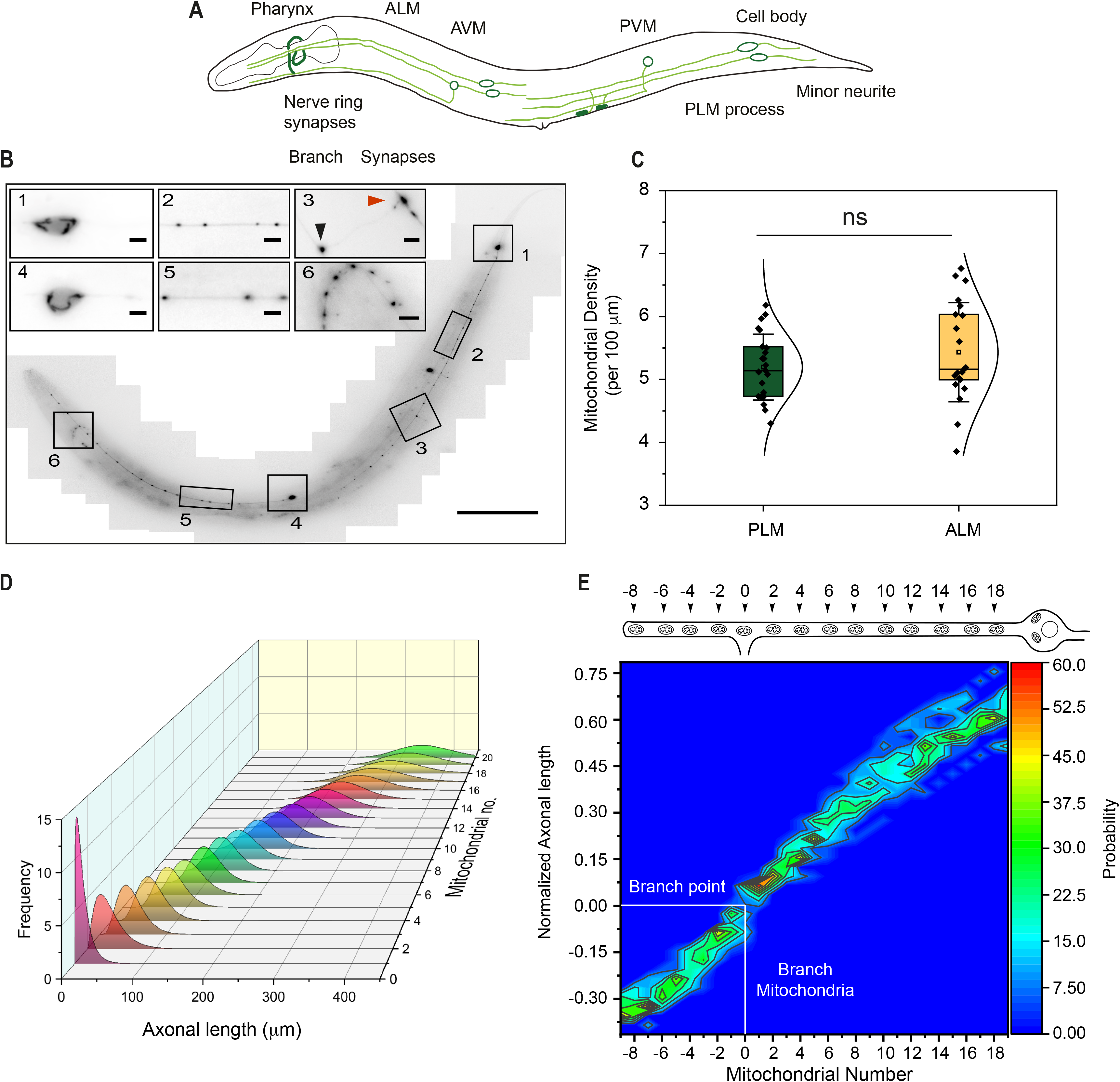
Mitochondrial positions in the TRN process are regulated. ****A****, Schematic *of C. elegans* TRNs. ****B****, Representative image of a 1d adult *C. elegans* expressing mitochondrially targeted GFP (*jsIs609*) imaged under a widefield microscope and stitched using ImageJ. Inset: Zoomed regions of the boxes shown in (****A****). Region1. PLM cell body; 2. Proximal PLM; 3. Branch point (Black arrow) and synapse (Red arrow); 4. ALM cell body; 5. Proximal ALM; 6. Nerve ring. ****C****, Distribution of mitochondrial density in 1d adult (Number of mitochondria per 100 micrometers axon length) along with a normal distribution fit. n >10 animals. ****D****, Distribution plot of every mitochondrial positions along PLM neuronal process. Gamma distribution fit of each mitochondrial positions from the cell body along the PLM is shown. n > 10 animals. Pink : First mitochondria from cell body. ****E****, Mitochondrial probability plot along the PLM neurons from 1d adult animals. Probability of finding a mitochondrion within ~10 μm neuronal process was calculated and presented as a heat map. Mitochondria at the branch point was assigned as ‘0’ mitochondria. n = 20 animals. Scale bar. 50 μm. Inset. 5 μm.

The observed constant mitochondrial density during growth can arise either by random positioning or regulated positioning of the mitochondria along the neuronal. To test how mitochondria are positioned along the neuronal process, we plotted the mitochondrial position along the axon to understand its distribution along the neuronal process. Positioning of mitochondria along the neuronal process follow a normal distribution (Fig. 1*D*). First mitochondria in young adults (1day old, 1d adult hence after) is positioned at 12.21±2.18 μm (Mean±SD) distance from the cell body (referred as proximal part). We also find that the SD of mitochondrial position in proximal part (2^nd^-4^th^ mitochondria) is smaller compared to distal part (18-20^th^ mitochondria). This indicates the positioning of mitochondria in TRN is regulated and positioning in proximal region is more tightly regulated than distal region (Variance of 1^st^ mitochondrial position 109.69, Variance of 20^th^ mitochondrial position 3136.9, *** p<0.001).

Additionally, random positioning of mitochondria across the animal will have decreased and uniform probability throughout the process while mitochondria located at a regulated distance will show enriched probability at regions where most-likely a mitochondrion will be found across different animals. We sought to test this by constructing a mitochondrial probability map along the entire process length taking advantage of the observation that a mitochondrion is present at the PLM branch point in all animals. We fixed the branch point as position ‘0’ (Fig. 1*D*) and measure the distance of every mitochondrion from this point. Since the length of the neuronal process is variable across animals, we normalized these distances to total neuronal length (normalized neuronal length). We then plotted our data as a histogram using a bin size of 0.03 (that roughly translate to ~10 μm neuronal length in un-normalized neuron) (Supplementary Fig. S2). Histograms of normalized distances from the branch point were further converted to a probability map of finding a mitochondrion at a given distance from the branch point. Figure 1*D* showed that, instead of a uniform probability heatmap, we observed locations with enriched probability along the entire length of the neuronal processes with respect to the branch point. Probability of finding a given mitochondrion along the neuronal process varies between 20-60%. The probability of predicting the adjacent two mitochondria is the highest (>50%) for the mitochondria adjacent to the branch point mitochondria. This probability falls over the axonal length either towards the cell body or towards the end of the neuronal process. Mitochondria before the branch point have probability of ~60% while 11^th^ mitochondria from the branch point towards the cell body show only to 20% probability. This pattern holds true even when other mitochondria were chosen as reference (Supplementary Fig. S3). This data revealed that spatial positioning of each mitochondria along the TRNs are regulated such that a given mitochondria possibly determines the position of each adjacent mitochondria beside it.

### Mitochondrial distribution in TRNs is non-random

Mitochondrial distribution varies over neuronal types such as in chicken DRG neurons where it is reported to be uniform while in cerebellar axons in rodents *in vivo* is reported to be non-uniform (Miller, 2004; Ohno et al., 2011). To examine mitochondrial distribution through *C. elegans* TRN process, we simulated the mitochondrial positioning by using a random number array that randomly put mitochondria along a given axonal length while keeping the density ~5. Further, all adjacent inter-mitochondrial distances were measured from simulated neuron and compared them with the experimentally obtained inter-mitochondrial distances from 1d adult animals. After 100 simulations (for L1 and L2, while 50 simulations were used for L3 to 1d adult) all adjacent inter-mitochondrial intervals were calculated and plotted as a histogram (Fig. 2*A*). The histogram of inter-mitochondrial intervals between adjacent mitochondria shows an exponential decay in simulated neuron such that the largest number of mitochondria are placed <3 μm apart. However, in *C. elegans* TRNs the histogram of experimentally obtained adjacent inter-mitochondrial distances in both ALM and PLM shows a gamma distribution (with median inter-mitochondrial distance ~ 18 μm Fig. 2*B*, Supplementary Fig. S4). These data indicate that mitochondrial positioning in the TRNs is non-random and probably maintained by a regulatory mechanism such that only a few mitochondria (~2-3%) are placed <3 μm apart. To see, if mitochondria positioning is regulated beyond adjacent mitochondria, we repeated the mitochondria probability map of every third mitochondria. We observed that in 1d adult experimentally obtained probability of finding every third mitochondria (n+2 distances) is higher (~50%) for some mitochondrial position (particularly, the probability between 3^rd^ and 5^th^, ~43% and 5^th^ and 7^th^ ~52%) (Fig. 2*C*). In contrast, when we randomly placed mitochondria in neuronal processes (e.g. simulation), the probability of finding the third mitochondria is strongly decreased (the probability between 3^rd^ and 5^th^ is ~26%, and 5^th^ and 7^th^ is only ~21%) (Fig. 2*C*) indicating that mitochondrial position is TRNs is non-random. This data also suggests that a given mitochondrion may be able to locally regulate the position of a few adjacent mitochondria.

**Figure 2.**
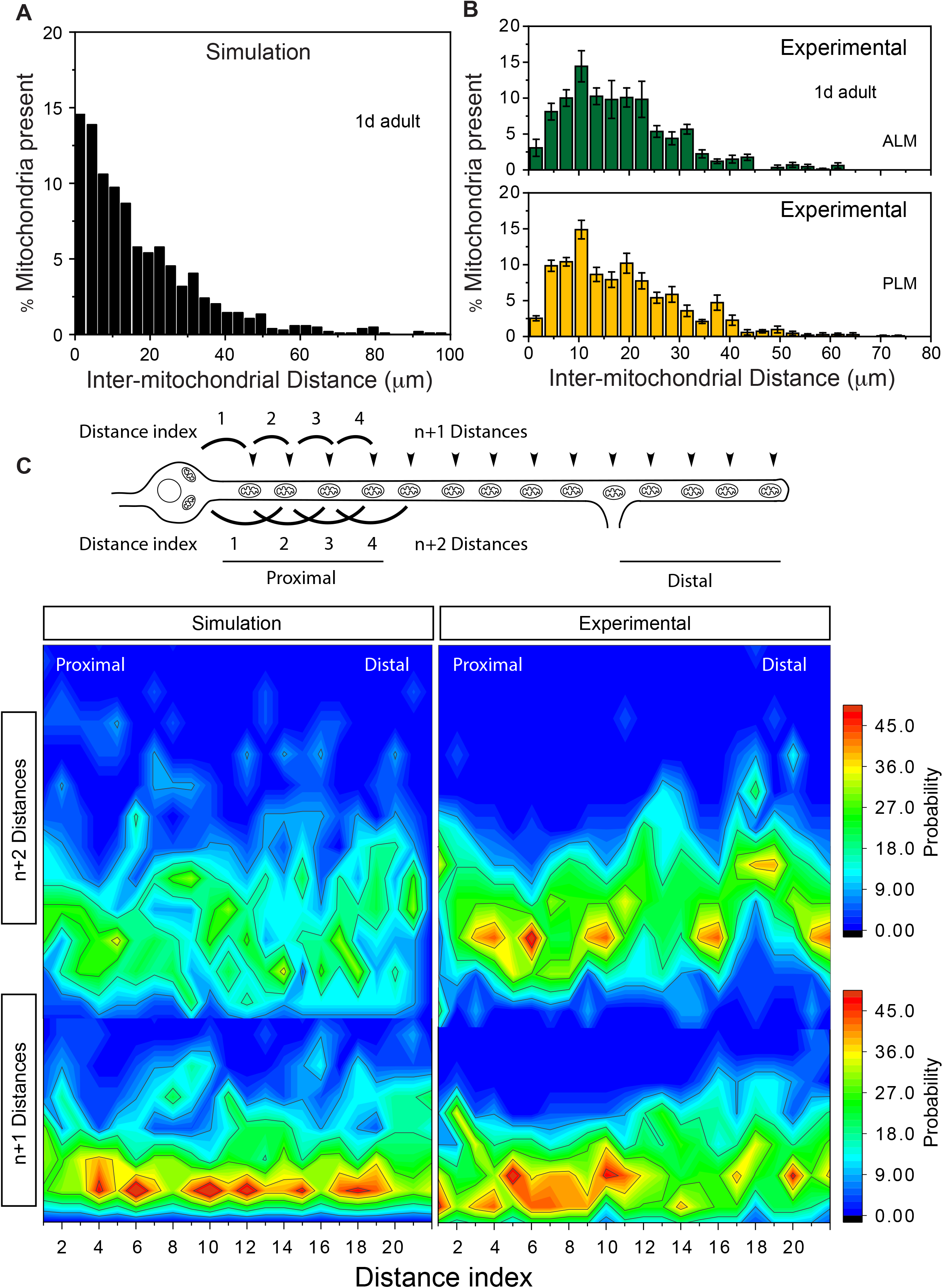
Mitochondrial positioning in the TRN process are non-random. ****A****, Monte-Carlo simulation of mitochondrial distribution. Mean neuronal length (395.66±47.43 μm) and mean density (5.32±0.84 per 100 μm) over 50 animals used for simulation and the distribution of inter-mitochondrial distances from 50 axons represented. ****B****, Distribution of experimental inter-mitochondrial distances from 1d adult ALM (green) and PLM (yellow) is shown. n>20 animals. ****C****, Contour plot of adjacent mitochondrial distances (n+1 distances) and distance between every third mitochondria (n+2 distances) from simulation (left) and experimentally obtained from PLM neurons (right) in 1d adult. n> 20 animals.

### Mitochondrial distribution remains constant throughout the development

To examine if mitochondrial distribution was developmentally regulated, we plotted histograms of all inter-mitochondrial distance and compared them with corresponding stage specific simulations (Fig. 3). Inter-mitochondrial distances from simulations corresponding to L1 to L4 larval stages revealed an exponential distribution similar to 1d adult. This indicates that unregulated, random positioning of mitochondria do no change the mitochondrial distribution as animal matures to 1d adult. However, histograms of measured inter-mitochondrial distances from all larval stages showed a gamma distribution similar to that seen in 1d adult (Supplementary Fig. S5) indicating the regulation that was observed in 1d adult is maintained throughout the developmental stages. Interestingly, the L1 experimental inter-mitochondrial distances shows a gamma distribution (Shape factor = 1.41, Scale factor = 14.21 for L1) closer to the exponential distribution seen in the corresponding simulation obtained for 1d adult (Simulation : Shape factor = 0.99, Scale factor = 18.14, Experimental : Shape factor = 2.67, Scale factor = 7.6 for 1d adult) or L1. In L1, a larger percentage of mitochondria are placed < 3 μm apart (~11.39%) that differs significantly from 1d adult (~3.37%) (Fig. 3, Supplementary Fig. S6*A, B*). This change in fewer mitochondria being placed < 3 μm apart occurs from the L2 developmental stage (Fig. 3, Supplementary Fig. S6*A, B*). There is no significant change in the fraction of mitochondria that are < 3 μm apart from L2 to 1d adult. This was also seen in cumulative probability plots of inter-mitochondrial distances from L1 to 1d adult (Supplementary Fig. S6*C, D*). Taken together, these data indicate that inter-mitochondrial intervals in TRNs are developmentally regulated and this regulation initiates from L2 developmental stage.

**Figure 3.**
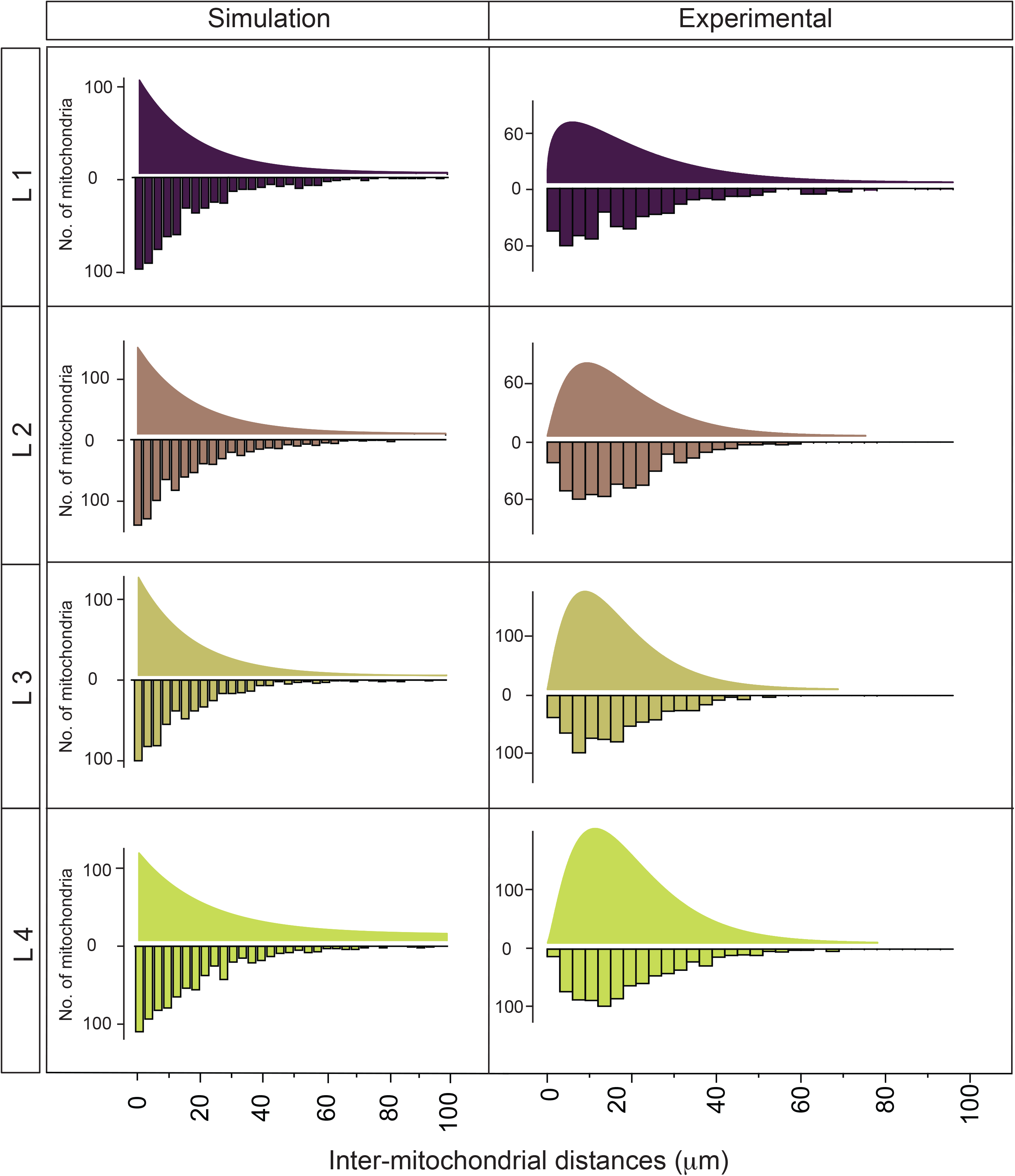
Mitochondrial distribution in the TRNs is non-random. Comparison of simulated (from > 50 animals with keeping density same) (Left) and experimentally obtained inter-mitochondrial distances (Right). 100 simulations were used for L1 and L2 stage to keep total mitochondrial number similar. Horizontal violin plot of raw distribution from all the distances shown in the bottom half while the fitted distribution of exponential (for simulation) and gamma function (Experimentally obtained) shown in the upper half. n~ 50 animals.

### Mitochondrial distribution is regulated by axonal transport

Previously, we showed that mitochondrial density in TRNs processes is reduced in mutants of the anterograde motor proteins while density remains unchanged in mutants of the retrograde motor dynein or in the mitochondrial trafficking adapter Miro-1 (Sure et al., 2018). To test if mitochondrial distribution observed in Figure 1 is dependent on anterograde and retrograde transport, we measured adjacent inter-mitochondrial intervals in mutants of Kinesin heavy chain *unc-116(e2310)* and dynein heavy chain *dhc-1(js319)*. In *unc-116* animals, despite reduced mitochondrial density (Fig. 4*A*), mitochondria are present throughout the neuronal process in the ALM and occasionally present at synapses (Supplementary Fig. S7*A*). The cumulative probability plot of inter-mitochondrial intervals shows significantly increased inter-mitochondrial distances in *khc-1* and similar inter-mitochondrial distances in *dhc-1* (Fig. 4*B*). This data indicates, mitochondrial distribution is strongly dependent on anterograde transport while retrograde transport may not play a major role.

**Figure 4.**
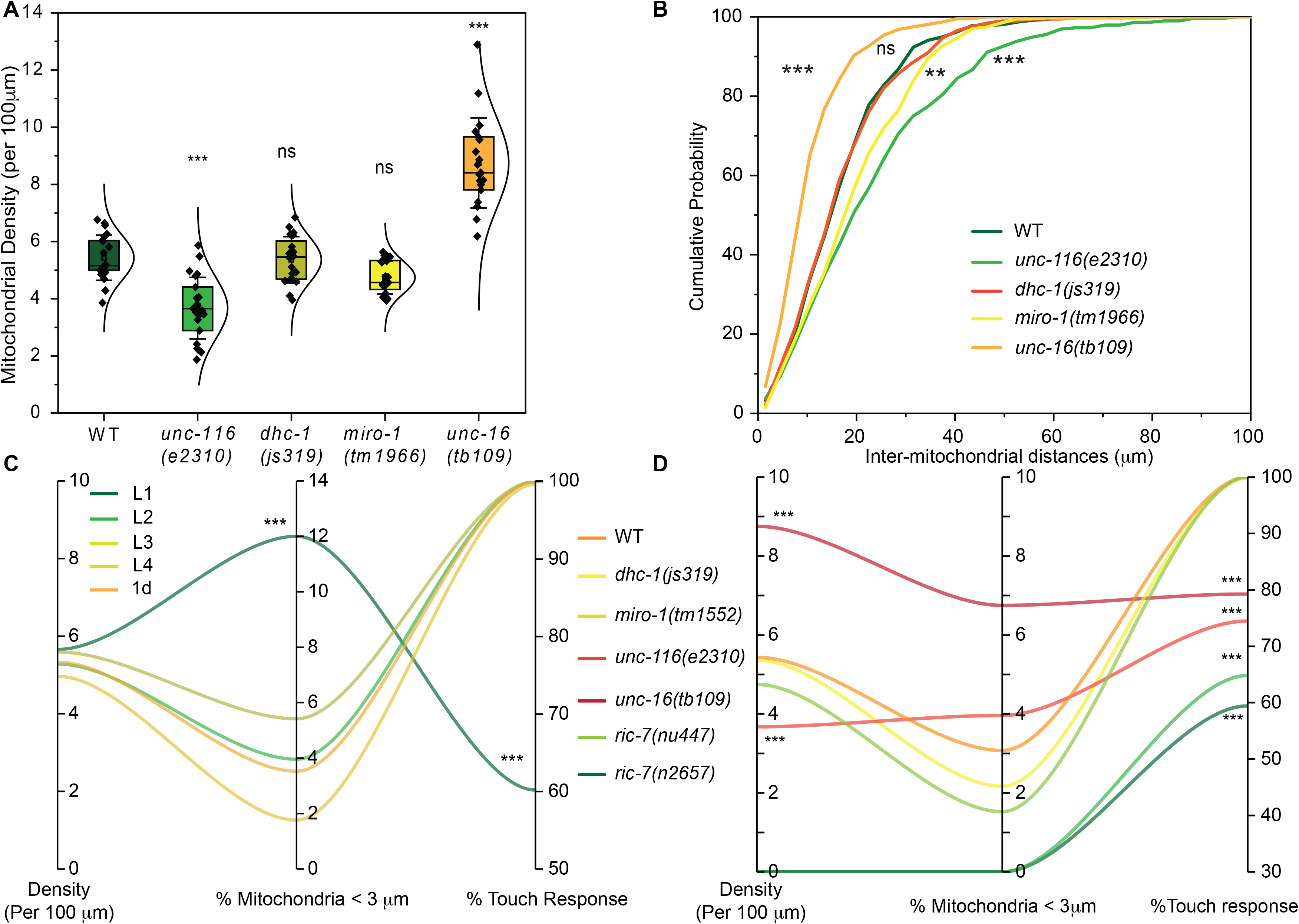
Mitochondrial distribution is regulated by axonal transport. Mitochondrial density (****A****) and distribution of inter-mitochondrial distances (****B****) in ALM neurons of motor mutants (*unc-116, dhc-1*) and its adaptors (*miro-1 and unc-16*). We did not see any mitochondria in the neuronal process in *ric-7(lf)* animals. n > 10 animals. ****C****, Parallel plot of mitochondrial density, percentage of mitochondria present μ3 μm apart and percentage touch response from 10 repetitive touches at different developmental stages in wild type (*jsIs609)* (****C****) and in motor and motor adaptor mutants (****D****). For mitochondrial density, percentage of mitochondria present μ3 μm apart n>10 animals, for % touch responsiveness 1d adult animals were chosen. n > 30 animals. ** p <0.01, *** p < 0.001, ns = non-significant.

The mitochondrial adaptor Miro is known to bind the motors Kinesin and Dynein to transport mitochondria bi-directionally (Morlino et al., 2014; Saotome et al., 2008). In *miro-1(tm1966)* mutants, we did not observe a significant change in their positioning (Supplementary Fig.S7*B*), inter-mitochondrial distances or density (Fig. 4*A, B*) (Sure et al., 2018). This is consistent with our earlier observations that in TRNs mitochondrial density is independent of Miro (Fig. 4*B*). Mutants in the *C. elegans* homolog of mammalian JIP3, *unc-16*, show increased mitochondrial density in the TRN process (Sure et al., 2018). We observed *unc-16* animals with increased mitochondrial density decreased inter-mitochondrial intervals with a significantly increased percentage of mitochondria <3 μm apart (Fig. 4*B*). Together, these observations suggest that mitochondrial density possibly determine inter-mitochondrial distances and proteins that regulate anterograde trafficking can alter the proximity of two adjacent mitochondria.

### Touch responses depend on mitochondrial density and inter-mitochondrial distances

*C. elegans* TRNs are responsible for sensing gentle touch applied to the body of the worm and in turn contribute to the corresponding response (Chalfie et al., 1985). We wished to examine whether a change in mitochondrial density or a change in the fraction of adjacent mitochondria <3 μm apart affects touch sensation. Therefore, we examined touch responsiveness over all developmental stages and in mutants with altered density and adjacent inter-mitochondrial intervals.

Gentle touch responsiveness in wild type is lowest in L1 and is invariant L2 onwards (Supplementary Fig. S8*A, B*). The observation correlates well with the inter-mitochondrial distance distribution where the only change between L1 and other developmental stages is the fraction of adjacent mitochondria <3 μm and not the density. A representation of the correlation between touch responsiveness and distribution is readily apparent in the Parallel plot shown in Figure 4*C* (Supplementary Fig. S9*A, B*). These data suggest that gentle touch responsiveness may be influenced by how close adjacent mitochondria are placed along the TRNs.

We also used mutants to assess the role of mitochondria in touch responsiveness. In both *ric-7(lf)* alleles (*nu447* and *n2657*) where mitochondria are known to be absent from neuronal process (Rawson et al., 2014), touch responsiveness was significantly reduced (64.8±2.06% and 59.41±1.98% respectively Mean±SEM) compared to wildtype (100%, Supplementary Fig. S8*C*). In *unc-116(e2310)*, where both mitochondrial density and fraction of adjacent mitochondria <3 μm were reduced, touch responsiveness was also significantly decreased (74.47±2.19 Mean±SEM) (Supplementary Fig. S8C). By contrast, *dhc-1(js319)* and *miro-1(tm1966)* animals with normal mitochondrial densities and adjacent inter-mitochondrial intervals did not show any significant change in touch responsiveness (99.81±0.18% for *dhc-1* and 100% for *miro-1* respectively) when compared to WT (Supplementary Fig. S8*C*). *unc-16(tb109)* mutant showed significant increase in mitochondrial density and a reduction in adjacent mitochondrial intervals, also shows significantly reduced touch responsiveness (79.26±1.87% Mean±SEM) (Supplementary Fig. S8*C*). The dependence of touch responsiveness in some of these mutants on the presence of mitochondria, its density and adjacent inter-mitochondrial distances is apparent in the parallel plot (Fig. 4*D*, Supplementary Fig. S9*C, D*). These data suggest that both mitochondrial density and adjacent inter-mitochondrial distances along the TRN neuronal processes are important for touch responsiveness.

### TRN specific rescue of mitochondria can rescue Touch response

To test whether mitochondrial density or fraction of mitochondria present <3 μm are important for touch responsiveness, we took advantage *ric-7(lf)* mutants where mitochondria are absent from the neuronal process (Rawson et al., 2014; Zhao et al., 2018) (Fig. 5*A*). Artificially driving mitochondrial transport by targeted expression of kinesin heavy chain directly to mitochondria (Mitotruck or mTruck) (Rawson et al., 2014), restores mitochondria into the neuronal processes of *ric-7*. In *ric-7* worms expressing mTruck under the TRN specific *mec-4* promoter, mitochondria are found to be present throughout the neuronal process in both ALM and PLM including at synaptic regions (Fig. 5*A, B*). Expression of mTruck restores ~12 mitochondria (11.7±2.55, Mean±SD) in the TRN processes of *ric-7(lf)* mutants which is significantly lower than WT (24.4+2.69) animals or WT animals (20±4.53) expressing mTruck (Supplementary Fig. S10). This reduced number of mitochondria resulted in a significantly lower density per 100 μm (3.87±0.16, Mean±SEM)) in *ric-7(lf)* animals compared to WT (5.43±0.17) and WT animals expressing mTruck (4.48±0.22) (Fig. 5*C*). Additionally, we also observed that the proximal part (41.5±11.3 in the first 100 μm) of the neuronal processes has a greater fraction of mitochondria than the distal part (18.2±9.2 μm in the last 100 μm) (Fig. 5*A*, Supplementary Fig. S10). To test if restoration of TRNs alone in *ric-7;mTruck* is sufficient to rescue the touch responsiveness, we performed the gentle touch assay. We observed that artificially driving mitochondria in the TRN processes of *ric-7(lf)* mutants restores touch responsiveness to that seen in WT and WT animals expressing mTruck (Supplementary Fig. S11*A*). The Parallel plot shows that the behavior correlates with the presence of appropriately spaced mitochondria in the proximal regions of the TRN processes are sufficient to restore touch responsiveness (Fig. 5*D*, Supplementary Fig. S11*B, C*). Overall, this data indicates that in addition to mechanosensory channels, presence of mitochondria in the TRNs is important for gentle touch sensation. In addition, our data also suggests that rescuing mitochondria only in the six TRNs is sufficient to rescue defect in touch sensation observed in *ric-7* indicating TRN mitochondria play a regulatory role in mechanosensation.

**Figure 5.**
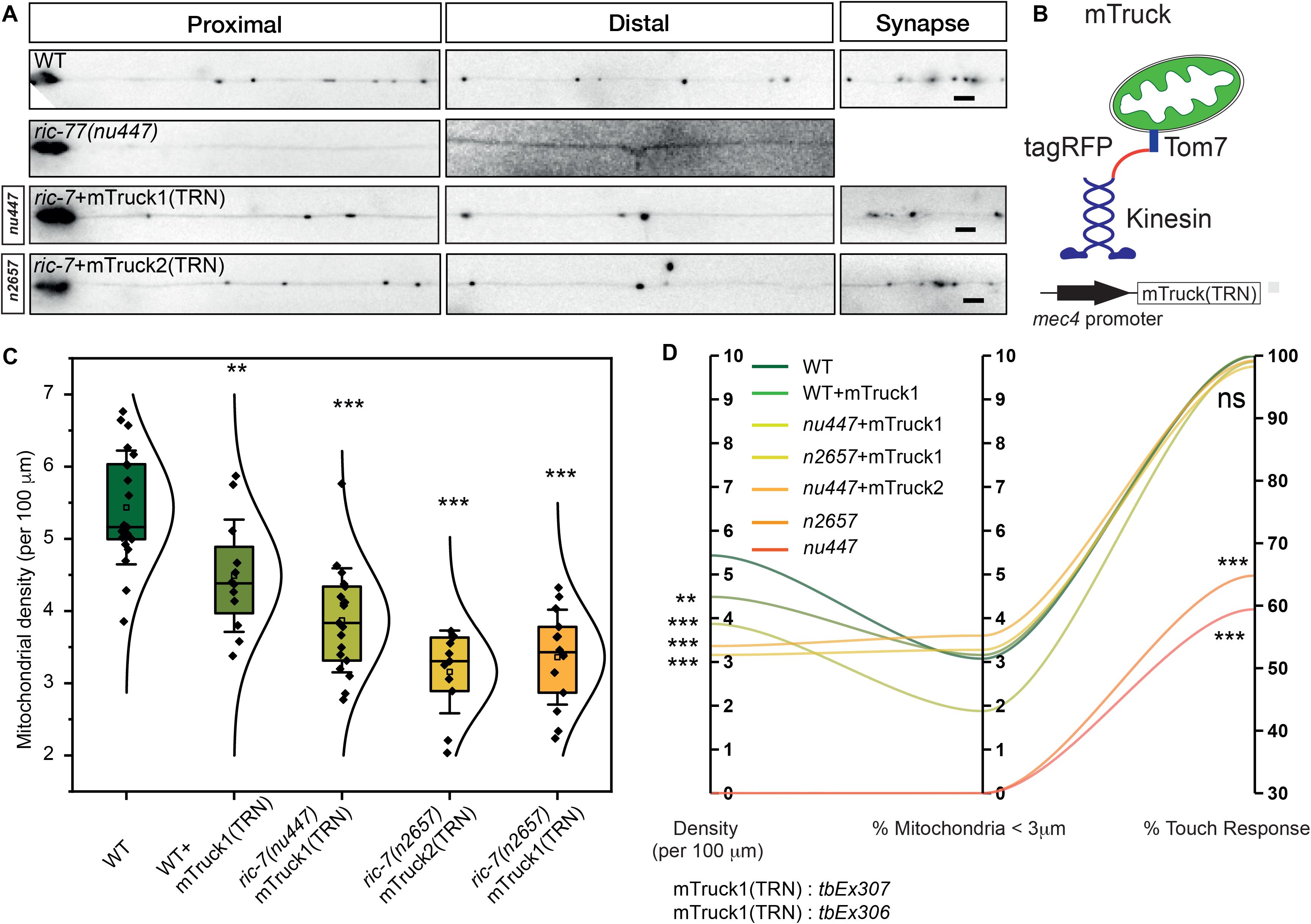
Mitochondrial presence is crucial for gentle touch responsiveness. ****A****, Positioning of mitochondria along the different segments of PLM neuron in WT, *ric-7(lf)* and *ric-7(lf)* mutant animals expressing TRN specific mitoTruck. ****B****, Schematic of TRN specific mitoTruck (mTruck). ****C****, Density of mitochondrial along the ALM neurons. n > 10 animals. ****D****, Parallel plot of mitochondrial density, percentage of mitochondria present μ3μm apart and percentage touch responsiveness from 10 repetitive touches in WT, *ric-7(nu447), ric-7(n2657)* and several *ric-7* alleles expressing mTruck(TRN). For mitochondrial density, percentage of mitochondria present μ3 μm apart n > 10 animals, for % touch response n > 30 animals. ** p <0.01, *** p < 0.001. Scale bar. 5 μm.

## Discussion

Results described here demonstrated that mitochondrial density and distribution in *C. elegans* TRNs are developmentally regulated. This regulation may arise during Larval stage 1 to Larval stage 2 transition and mitochondrial density and distribution are strongly dependent on anterograde motors and motor adapters. Additionally, this distribution is important for neuronal function as a change in both density and distribution alters gentle touch response of the animals. This report highlights the importance of mitochondrial positioning along the neurons *in vivo* and explores the role of mitochondria during neuronal communication in intact circuits.

### Mitochondrial positioning is non-random

The axonal positioning of mitochondria is important for the proper development, survival, and function of neurons (Iijima-Ando et al., 2012; Sun et al., 2013; Vaarmann et al., 2016). During *Drosophila* larval development from 1st instar to 3rd instar stage mitochondrial density in motor neurons remains constant and follows a linear relationship with axonal growth (O’Toole et al., 2008). While we and others showed that mitochondrial density in *C. elegans* TRNs remains constant over larval development and ageing (Morsci et al., 2016). We observed that each mitochondrion is normally distributed along the neuronal process and the positioning of adjacent mitochondria can be predicted. We also find that the median distance between adjacent mitochondria is ~18 μm. However, till now it remains unclear how two mitochondria maintain the distance between them during neuronal growth.

Earlier studies reported that the mitochondrial distribution along the DRG neurons from chicken embryo follows a uniform distribution (Miller, 2004). However, our results showed that in *C. elegans* mechanosensory neurons mitochondrial distribution is non-uniform and the inter-mitochondrial distances follows a gamma distribution which is significantly different than predicted exponential distribution by simulation. Moreover, comparing the differences in distribution at L1 stage with the rest of the developmental stages revealed major re-arrangement of mitochondrial positioning as animals mature to the 1d adult stage. We believe that at L1 stage mitochondria is transported from the cell body and positioned more randomly along the process. After L1/L2 molt, mitochondria are positioned in a more regulated manner which is responsible for the change in distribution as the animals reach adulthood.

### Mitochondrial density and distribution influences touch responses

The bi-directional transport of mitochondria mediated by molecular motors and adaptors are known to maintain mitochondrial density in vertebrates as well as invertebrates neurons (Saotome et al., 2008; Schwarz, 2013). However, in *C. elegans* TRNs mutations in retrograde motor Dynein and mitochondrial adaptor Miro did not significantly affect density or mitochondrial distribution (Sure et al., 2018). We observed that only mutation in Kinesin and kinesin associated proteins affecting anterograde transport, alters mitochondrial density and inter-mitochondrial distances. This further revealed that Kinesin mediated mitochondrial transport ensures proper mitochondrial density and distribution in *C. elegans* TRNs.

Density and localization of mitochondria are crucial for neuronal function (Billups and Forsythe, 2002; Iijima-Ando et al., 2012; Rangaraju et al., 2019). In cultured neurons alteration of mitochondrial density in neuronal process showed altered growth (Courchet et al., 2013), stability of neuronal process (López-Doménech et al., 2016) and synaptic communication (Vaccaro et al., 2017). In *C. elegans* motor neurons, mitochondrial trafficking mutants resulted in altered secretion from motor neurons to muscles and showed increased degeneration upon axotomy (Rawson et al., 2014; Zhao et al., 2018). Similarly, a recent report highlighted the importance of mitochondrial morphology (by fission and fission) affecting *C. elegans* locomotion and touch sensation (Byrne et al., 2019). Both studies indicated that the positioning or morphology of mitochondria directly influence behavior. Our results demonstrating mitochondria density and distribution contributes to touch responsiveness also reinforce these findings. Absence of mitochondria from neuronal process (*ric-7(lf)*), increase in proximity of adjacent mitochondria (e.g L1 stage or *unc-16)* or decrease in mitochondrial density (e.g. *unc-116*) significantly decreased touch response.

This modulation of touch response is strongly dependent on the presence of the mitochondria in TRNs as ectopically restoring mitochondria in only six TRNs can partially rescue the touch sensation in *ric-7(lf)*. Interestingly, ectopically restoring mitochondria in TRNs do not rescue density but restore touch response as observed in WT. This indicates only decrease in density may not be sufficient to alter touch responsiveness. Together, our data highlighted the importance of mitochondria for modulating the touch response from six TRNs. Inducing hypoxia like environment decreases touch response in the *C. elegans* by secreting several neuromodulators for signaling (Chen and Chalfie, 2014). Recently, it is reported that the absence of mitochondria from motor neurons of *C. elegans* can induce hypoxia like environment^33^. It is possible that mitochondria can play similar roles in TRNs and modulate touch responsiveness preventing the formation of hypoxia like environment.

## Acknowledgments

*jsIs609* was made by SPK in Dr. M. Nonet’s laboratory. We thank CGC (NIH Office of Research Infrastructure Programs (P40 OD010440) and Mitani Lab (National Bio-Resource Project of the MEXT, Japan) for strains. We thank Dr. Joshua Kaplan for mitoTruck plasmid. We acknowledge Drs. Gautam Menon and Varuni Prabhakar for useful discussion on mitochondria transport, image analysis and their suggestions on manuscript. SM was supported by Marie Skłodowska-Curie International Incoming Fellowship (Nos. 913033) and TIFR. Authors thank BITS (AA), TIFR (SH, AC), CSIR-HRDG (GPRS) and DBT postdoctoral fellowship (SM) for funding. This work is funded by DAE-PRISM 12-R&D-IMS-5.02.0202 (SPK and Gautam Menon), and HHMI-IECS grant number 55007425 (SPK).

## Materials and Methods

### Materials

60 mm plates were obtained from Praveen Scientific (New Delhi, IN), Bacto Agar and Peptone were obtained from BD bioscience (New Jersey, USA), NaCl was obtained from Hi-Media (Mumbai, IN), Cholesterol, CaCl2, MgSO4 and Sodium Azide was obtained from Sigma-Aldrich (St. Louis, Missouri, United States). For imaging cover slides and coverslips were obtained from Bluestar No.1 coverslips (Mumbai, IN).

### Worm maintenance and imaging

Worms were grown at 0°C on NGM agar media seeded with *E. coli* strain OP50 under standard laboratory conditions. Worms were transferred to a fresh plate at larval stage 4 and imaged at 1d adult. All the imaging was performed at 1d adult unless specified using 1-10 mM sodium azide (prepared in M9 buffer) as an anaesthetic. Animals were mounted in a coverslip and imaged under an upright Nikon E800 epifluorescence microscope (Tokyo, Japan) microscope or an inverted Olympus IX73 epifluorescence microscopes (Tokyo, Japan) equipped with either EvolutionQEi Monochrome camera or Photokinetics Evolve EMCCD camera respectively. Imaging was performed at 60× magnification and overlapping images were collected to reconstruct the entire neuronal process. The total neuronal lensgth was quantified using background GFP fluorescence in the TRN along with the number of mitochondria.

### Image analysis and simulation

All the image analysis was performed using ImageJ (NIH) with in-built plugins. For mitochondrial density per 100 μm axonal length, the total number of mitochondria and the total neuronal length was calculated. A cut-off of 2×2 pixel was used to identify smaller mitochondria, any fluorescent particle smaller than that was discarded. Inter-mitochondrial distances were calculated from the measured distances between the mid-point of two adjacent mitochondria. Additionally, we also consider the distance from the cell body and first mitochondria as well as the distance from last mitochondria and the end of the neuronal process. All the inter-mitochondrial distances then plotted as a histogram with 3μm bin size and further converted into cumulative probability map for statistical comparisons.

For distribution plot of mitochondria, mitochondrial location from cell body was determined across different animals and plotted as a violin plot with 10 μm bin. The mitochondria positions were tested for normality and a normal distribution fit of the mitochondrial histogram was performed in Origin2020b. The resulting normal distribution of each mitochondria was plotted as a 3D plot including mitochondrial number from cell body, axonal length and frequency of distribution.

For mitochondrial probability map of PLM neuron, overlapping images were stitched and neuron was straightened. X and Y co-ordinate of each mitochondrion were measured by ‘Find Maxima’ plugin and saved. Next, the co-ordinate of the mitochondria of the branch is marked as ‘0’ and all the mitochondrial co-ordinate was normalized with the respect it by subtracting its original co-ordinate. This normalization makes the mitochondria towards cell body having positive co-ordinate while mitochondrial co-ordinate after the branch has negative coordinates. Since each neuronal length is different, we further divide these co-ordinates with a total length of the neuron and plotted co-ordinate of each mitochondrion as a histogram (with ~10 μm bin) and from the histogram a percentage probability map (% of mitochondria at a given bin of ~ 10 μm) was constructed.

Similar method was followed for calculation of probability of adjacent mitochondria or every third mitochondria. Distance from cell body to first mitochondria is chosen as distance index 1 and then n+1 distances was computed in Origin2020b software (distance index of 2 = first mitochondria from cell body-second mitochondria from cell body and continued). For n+2 distances, distance of second mitochondria from the cell body was chosen as distance index 1 and then continued (distance index of 2 = first mitochondria from cell body-third mitochondria from cell body and continued).

Monte-Carlo simulation was performed in Matlab/octave online using measured neuronal length and mitochondrial density at a given developmental stage. Typically, at 1d adult stage, ~400 μm neuronal length was considered and, in this length, ~20 mitochondria were added randomly. We use 2×SD of neuronal length to vary neuronal length across animal and a random number array is used to vary the number of mitochondria (number of mitochondria also varied by considering 2×SD of mitochondrial density per 100 μm) along the given length. The inter-mitochondrial distances from randomly placed mitochondria were measured and plotted as a histogram of 3μm bin. For the L1-L2 stage, the simulation was performed using 100 animals while from L3-1d adult 50 animals were considered for simulation. The Matlab code has been provided in the supporting info.

### Gentle touch assay

*Plate preparation*: Plates poured 4 days before the assay and stored at 4°C were used to maintain the consistency in the water content of the plates across all the tests. Plates were seeded with 400 ml of an overnight grown culture of *E.coli OP50* (O.D. ~0.6) one day before the assay and stored at 4°C. *Gentle touch assay*: Animals in the mid-L4 stage were transferred on plates prepared for the assay and allowed to age for 5-6 hours until they enter the adult stage. The assay was performed in the temperature range of 20-25°C and Humidity varied from 50-70%. Gentle touch stimulation was provided with the help an eyelash attached to a stick. Animals were touched alternatively on the head between the pharynx and the vulva (anterior touch), and on the tail between the vulva and anus (posterior touch). A response was counted if animals moved backwards, accelerated (in case they are already moving backwards) or stopped moving when touched anteriorly. Similarly, a response to posterior touch was counted when animals moved forward (in case they are moving backwards), accelerated (in case they are already moving forward) or stopped moving. Animals were not touched near the vulval region to avoid an omega turn behaviour. The responses were counted using a cell counter wherein, a positive response was counted as 1 and a negative response was counted as 0. After repeated stimulation, the animal was removed from the plate to avoid assaying it again. The interstimulus interval (ISI) between an anterior and a posterior touch was maintained at 1 second using a counter which beeped after every second. The interval may extend in a few instances where the animals moved.

### Cloning and microinjections

MitoTruck(TRN) was prepared from *unc29p::unc-116::tagRFP::tom7* plasmid obtained from Kaplan lab (Dept. of Mol. Biol., Harvard University). Briefly, *unc-116::tagRFP::tom7* including *unc-54* 3’-UTR was PCR amplified using Phusion polymerase (NEB, Ipswich, MA, USA) from *unc29p::unc-116::tagRFP::tom7* using following primers, FP: 5’-AGCAAGGCTAGCCAAGACAAGTTTGTAC-3’ and RP: 5’-ACTCACGGGCCCTAGTGGGCAGATCTT-3’ and cloned between Nhe-1 and Apa1 (NEB, Ipswich, MA, USA) sites in *mec4p::Lamp-1::GFP*.

Plasmids were purified using Macherey-Nagel NucleoSpin Plasmid purification kit (Düren, Germany) and subjected to Ethanol precipitation before using. A cocktail of three plasmids, 20ng/ml *mec4p::unc-116::tagRFP::tom7*, 50 ng/ml and *ttxp::*RFP, 130 ng/ml PBluscript SK-was prepared and microinjected into 1d adult N2 stain using Eppendorf FemtoJet microinjector (Hamburg, Germany) fitted in Olympus IX53 (Tokyo, Japan).

### Statistical tests

All the data was plotted using OriginPro 2020 (Origin Lab, Northampton, MA, USA) and figures are prepared using Adobe Illustrator (Adobe Corporation, San Jose, CA, USA). For mitochondrial density, normality was calculated using Kolmogorov-Smirnov test and statistical significance was calculated either two-sample student’s t-test with Welch’s correction (for comparison between two conditions) or One-Way ANOVA with Bonferroni Post Hoc test (For >2 conditions). For inter-mitochondrial distance comparison, a nonparametric Kolmogorov–Smirnov test (KS test) test was performed. For gentle touch response, a non-parametric Kruskal–Wallis one-way ANOVA was used. p values *<0.05, **<0.01 and ***<0.001 was used for significance. p values >0.05 was considered non-significant.

### Genotypes used in this study

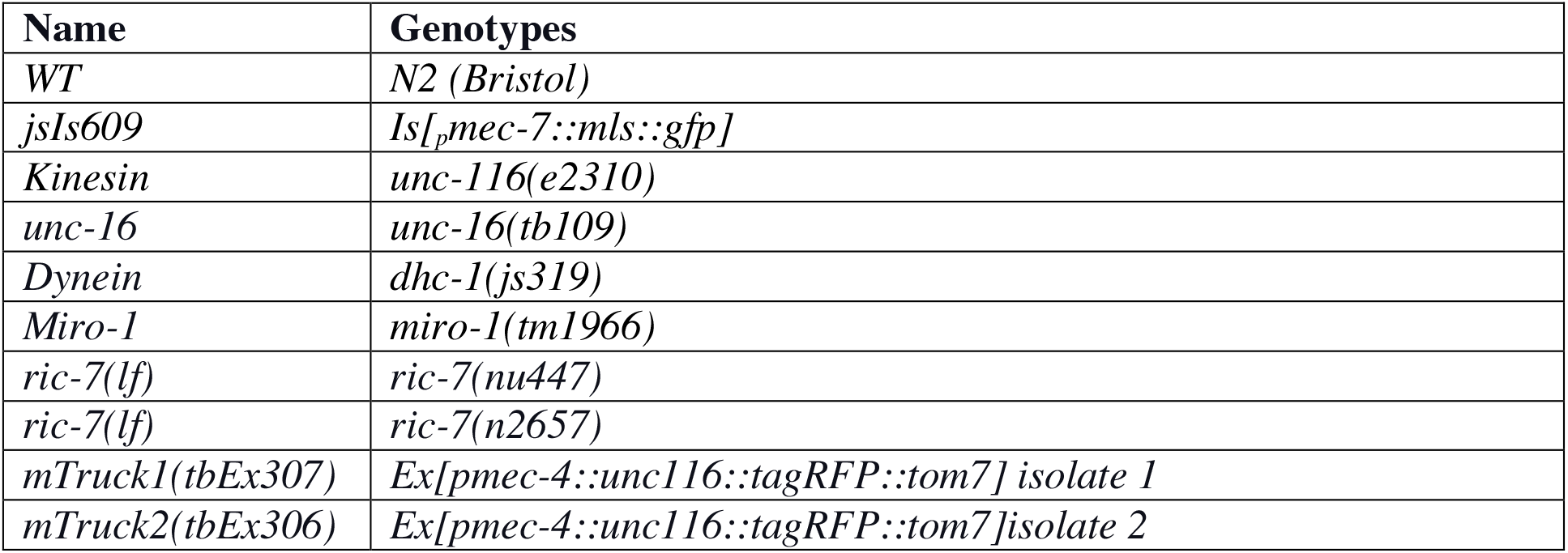

