## Supplemental information for "Regulated distribution of mitochondria in touch receptor neurons of *C. elegans* influences touch response"

### Supplementary Figure 1.

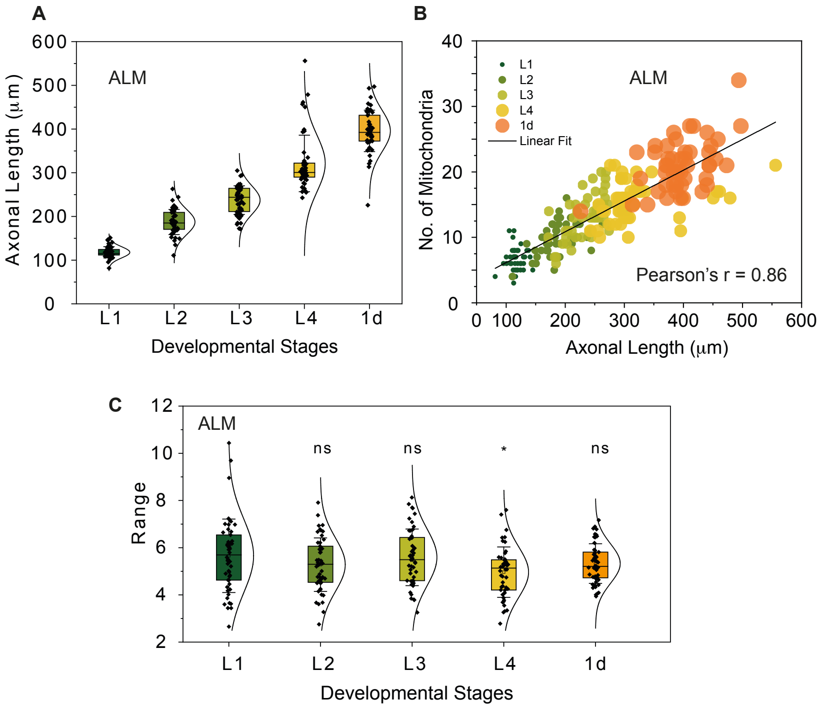


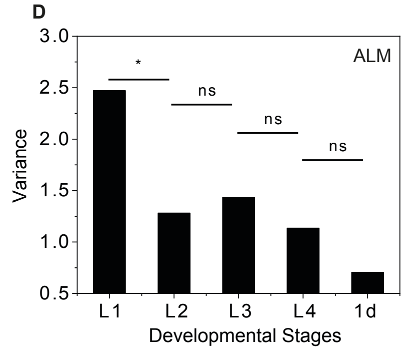


Table 1

| ALM | | PLM | |
|---|---|---|---|
| Comparison | F-Test | Comparison | F-Test |
| L1-L2 | 0.02821 | L1-L2 | 0.00832 |
| L1-L3 | 0.07267 | L1-L3 | 8.82E-04 |
| L1-L4 | 0.01023 | L1-L4 | 0.00228 |
| L1-1d | 3.15E-05 | L1-1d | 1.35E-04 |
| L2-1d | 0.0395 | L2-1d | 0.21615 |
| L3-1d | 0.01487 | L3-1d | 0.60141 |
| L4-1d | 0.10275 | L4-1d | 0.40312 |

**Supplementary Fig. S1.** ***A***, Linear increase of axonal length of ALM process over developmental stages. n ~ 50 animals. ***B***, Scatter plot of axonal length with respect to mitochondrial number in the PLM axonal process over different developmental stages. n ~ 50 animals. *C*, Violin plot of mitochondrial density from ALM and PLM neurons across all the developmental stages. L1: Green, L2: Light Green, L3: Light Yellow, L4: Orange, 1d adult: Red. n ~ 50 animals. ***D***, Bar graph of variance calculated from mitochondrial density in ALM process over different developmental stages. n ~ 50 animals. Table 1: Statistical test for variances observed in (***D***). ns = non-significant. * $p < 0.05$. **$p<0.01$, *** $p<0.001$. ns = non-significant.

**Supplementary Fig. S2.**

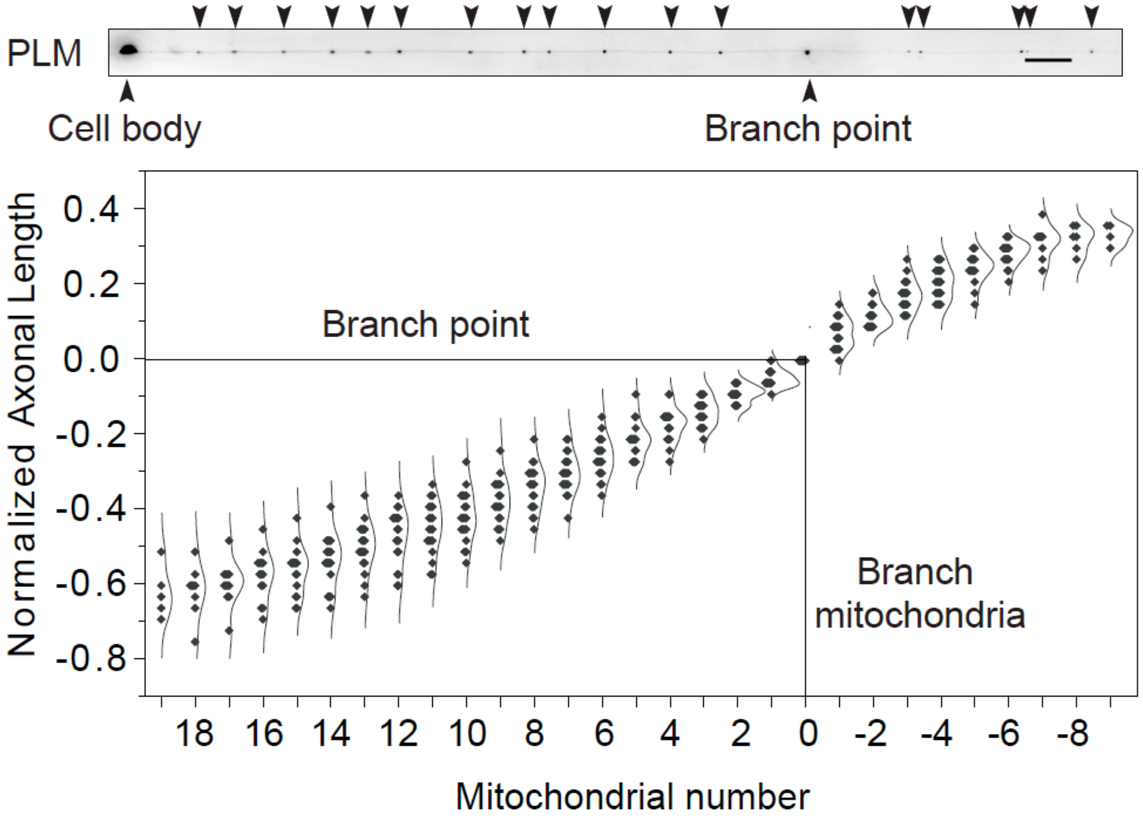


**Supplementary Fig. S2.** Histograms of mitochondrial positioning in the axonal process of PLM neurons. Branch mitochondria is assigned '0' and relative length of each mitochondria from branch point is calculated plotted as a violin plot. n ~ 20 animals.

**Supplementary Fig. S3.**

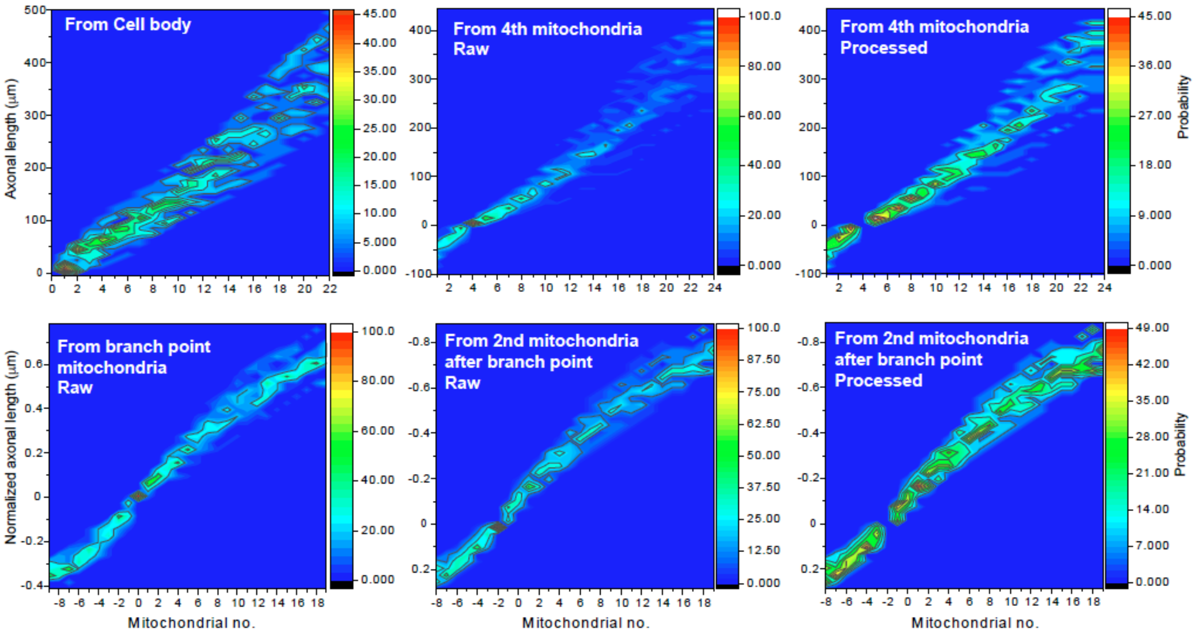


**Supplementary Fig. S3.** Probability plot of mitochondria located at different regions of the PLM neuronal processes. Raw: Probability plot of mitochondria chosen as constant (Note: probability of 100 % due to alignment). Processed: Probability plot of mitochondria chosen was omitted to enhance the visualization of the other mitochondrial positions.

### Supplementary Fig. S4.

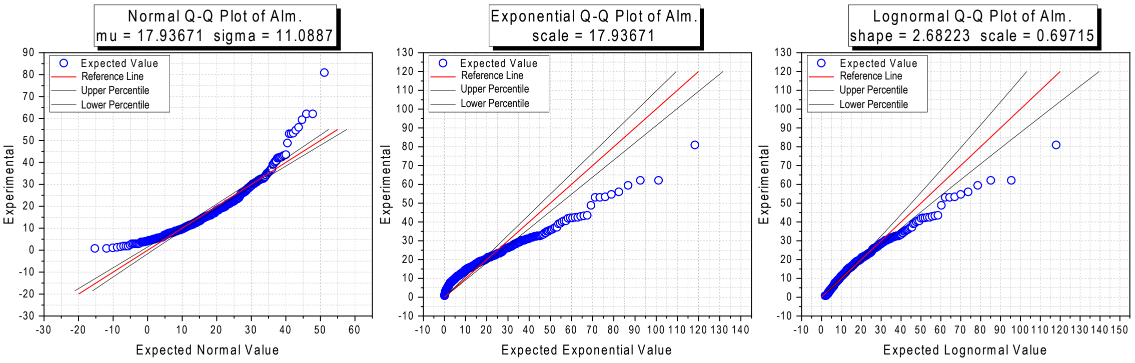


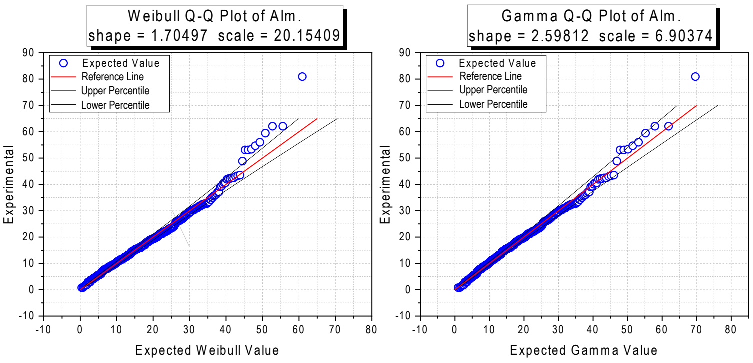


| Distribution | Goodness of Fit tests | Statistics | P-value | Decision at level(5%) |
|---|---|---|---|---|
| Normal | K-S modified test | 0.08966 | <=0.01 | Reject Normal |
| | A-D test | 7.64386 | 1.13E-18 | Reject Normal |
| Lognormal | K-S modified test | 0.07374 | <=0.01 | Reject Lognormal |
| | A-D test | 3.99208 | 5.98E-10 | Reject Lognormal |
| Weibull | K-S modified test | 0.04401 | >0.1 | Can't reject Weibull |
| | A-D test | 1.09468 | <0.01 | Reject Weibull |
| Exponential | K-S modified test | 0.20855 | <=0.01 | Reject Exponential |
| | A-D test | 34.40169 | 0 | Reject Exponential |
| Gamma | K-S modified test | 0.03095 | >0.25 | Can't reject Gamma |
| | A-D test | 0.50705 | 0.21886 | Can't reject Gamma |

**Supplementary Fig. S4.** Q-Q plot of inter-mitochondrial distances observed in 1d adult. Inter-mitochondrial distances were fitted with Gamma, Exponential, Normal or Lognormal distributions. Note, only gamma distribution can fit the distribution indicating, distribution of inter-mitochondrial distances follows a gamma distribution.

**Supplementary Fig. S5.**

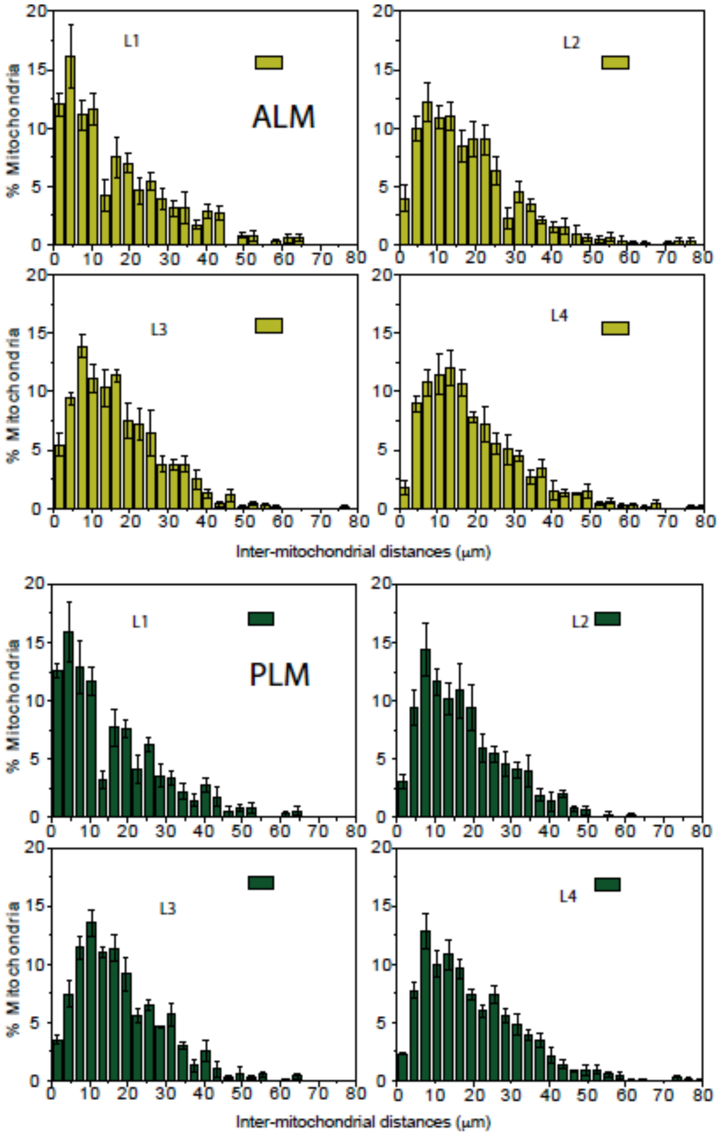


**Supplementary Fig. S5.** Distribution of inter-mitochondrial distances at different developmental stages as shown in Fig. 3. Percentage mitochondria in each bin is shown as Mean±SEM from both ALM and PLM neuronal processes. n ~ 50 animals.

**Supplementary Fig. S6.**

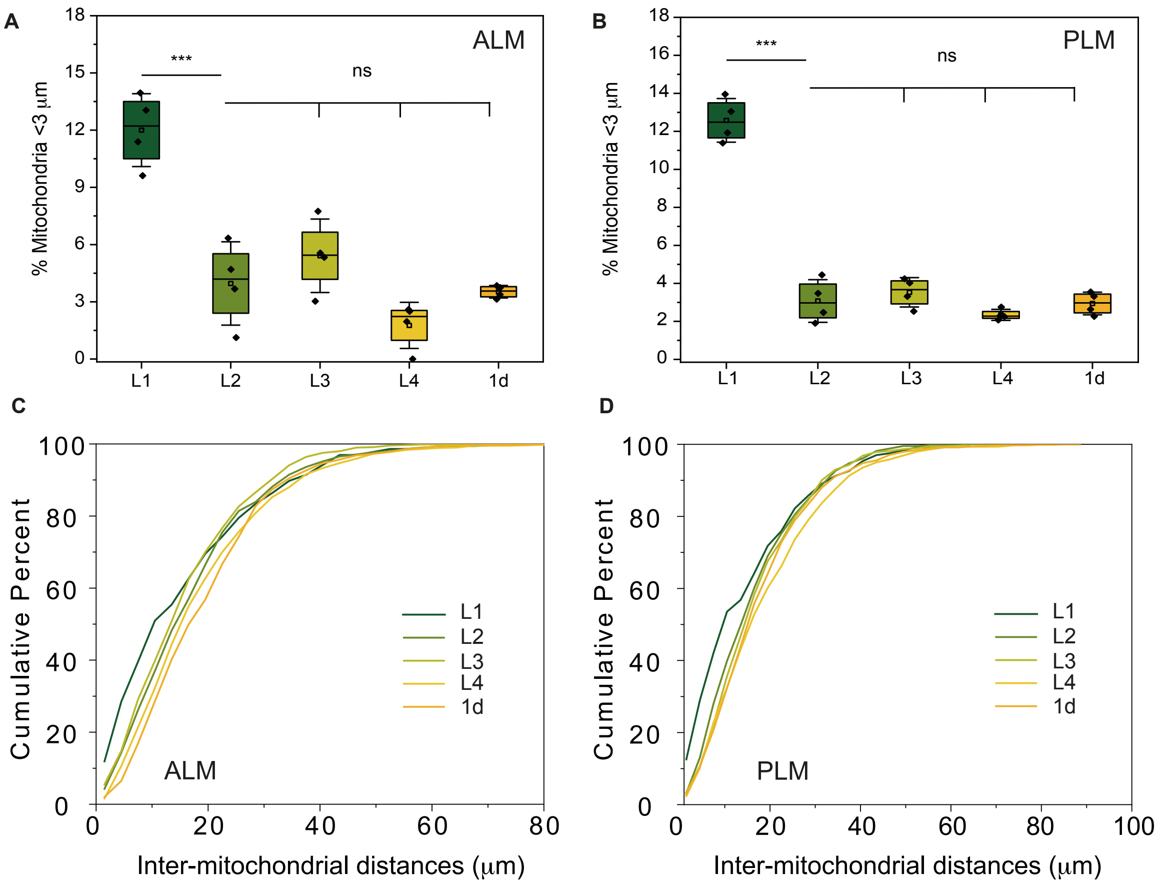


**Supplementary Fig. S6.** Mean percentage of adjacent mitochondria present <3µm apart across different developmental stages from ALM (***A***) and PLM (***B***) neuronal processes. n ~ 50 animals. ***p<0.001. ***C-D***, Cumulative probability plot of inter-mitochondrial distances across different developmental stages. n~50 animals.

**Supplementary Fig. S7*A*.**

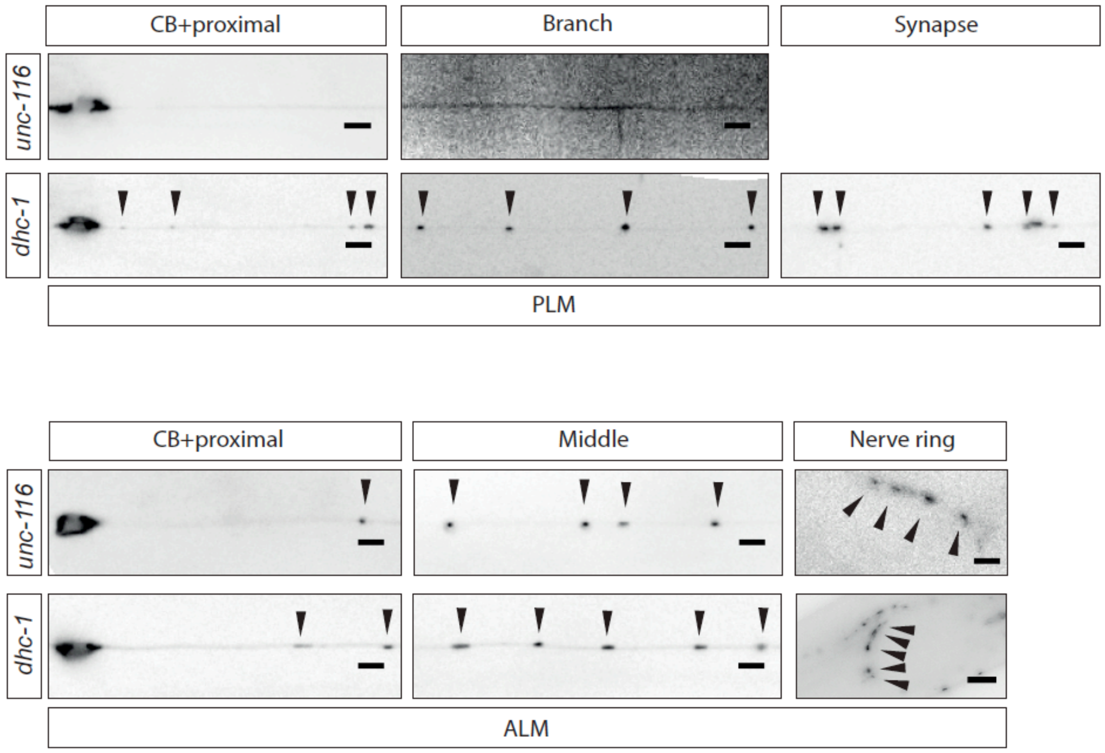


**Supplementary Fig. S7*A*.** Mitochondrial positioning along different segments of PLM and ALM neurons in Kinesin (*unc-116(e2310)*) and Dynein (*dhc-1(js319)*) mutants. Arrows indicate mitochondria. Scale bar. 5 µm, Scale bar. Nerve ring in *dhc-1*. 10 µm.

**Supplementary Fig. S7*B*.**

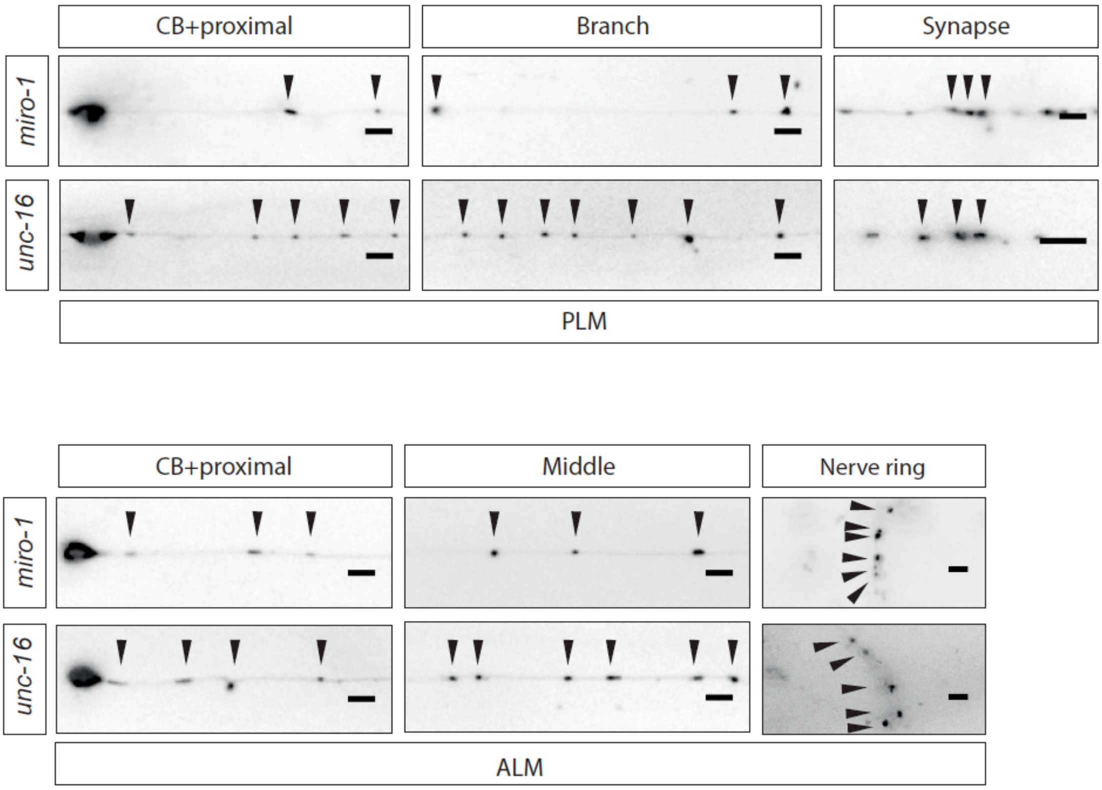


**Supplementary Fig. S7*B*.** Mitochondrial positioning along different segments of PLM and ALM neurons in motor adaptors *miro-1(tm1966)*) and *unc-16*(*tb109)*) mutants. Arrows indicate mitochondria. Scale bar. 5 µm

**Supplementary Fig. S8.**

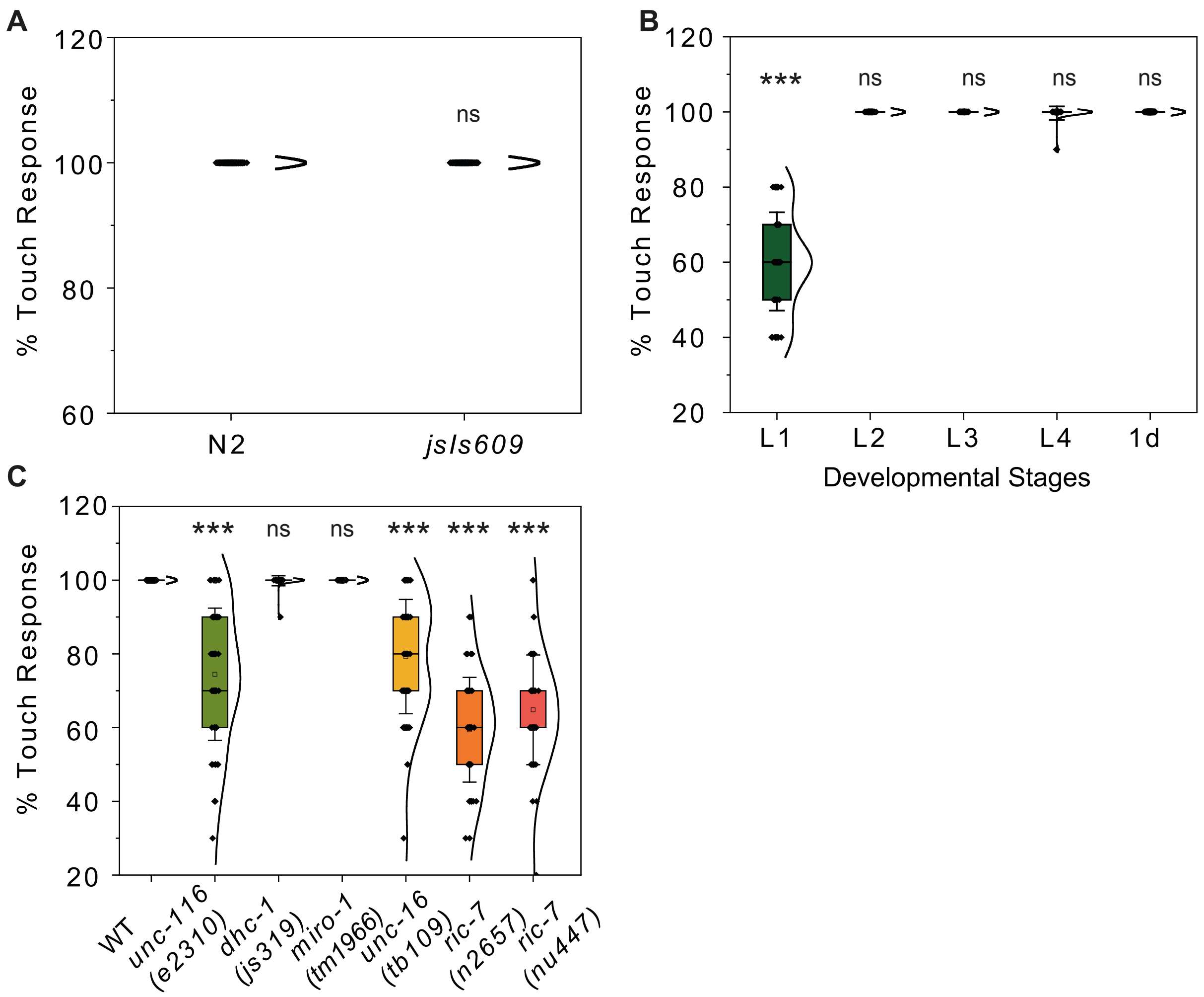


**Supplementary Fig. 8.** ***A***, Violin plot of percentage touch response in N2 and animals expressing MLS-tagged GFP (*jsIs609,* hence referred as WT). n > 50 animals. ***B***, Violin plot of percentage touch response in WT animals across different developmental stages. n > 30 animals. ***C***, Violin plot of percentage touch response in WT, motor mutants (*unc-116* and *dhc-1*), mutants of motor adaptors (*miro-1* and *unc-16*) and mutant of cytoplasmic protein aiding axonal transport of mitochondria (*ric-7*). n >30 animals. *** $p < 0.001$, ns = non-significant.

**Supplementary Fig. S9.**

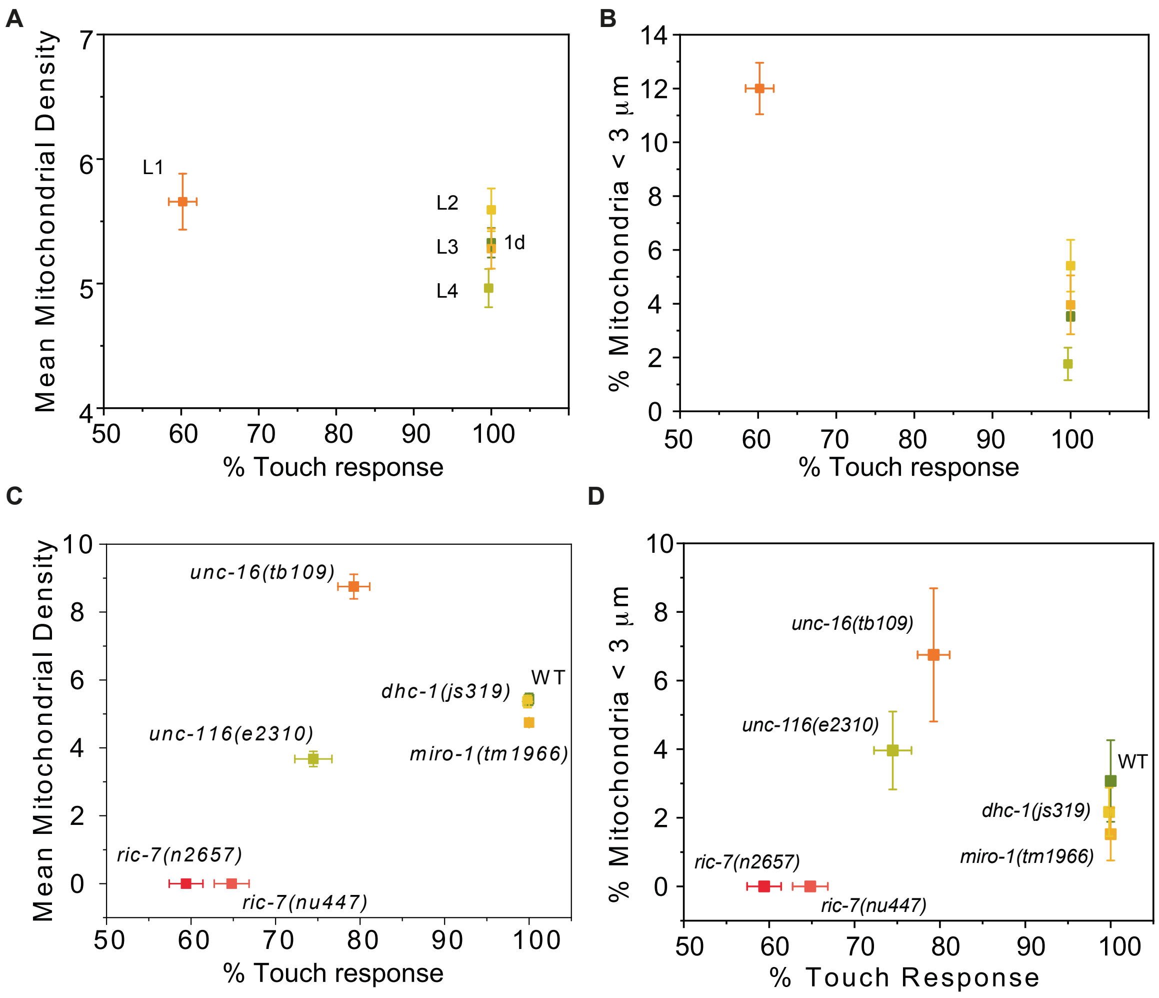


**Supplementary Fig. S9.** ***A-B***, 2D scatter plot of gentle touch response with respect to mean mitochondrial density and percentage of mitochondrial present <3 µm apart across developmental stages. ***C-D***, 2D scatter plot of gentle touch response with respect to mean mitochondrial density and percentage of mitochondrial present <3 µm apart in motor, motor adaptor mutants. For mitochondrial density, percentage of mitochondria present <3µm apart n > 10 animals, for % touch response n > 30 animals.

**Supplementary Fig. S10.**

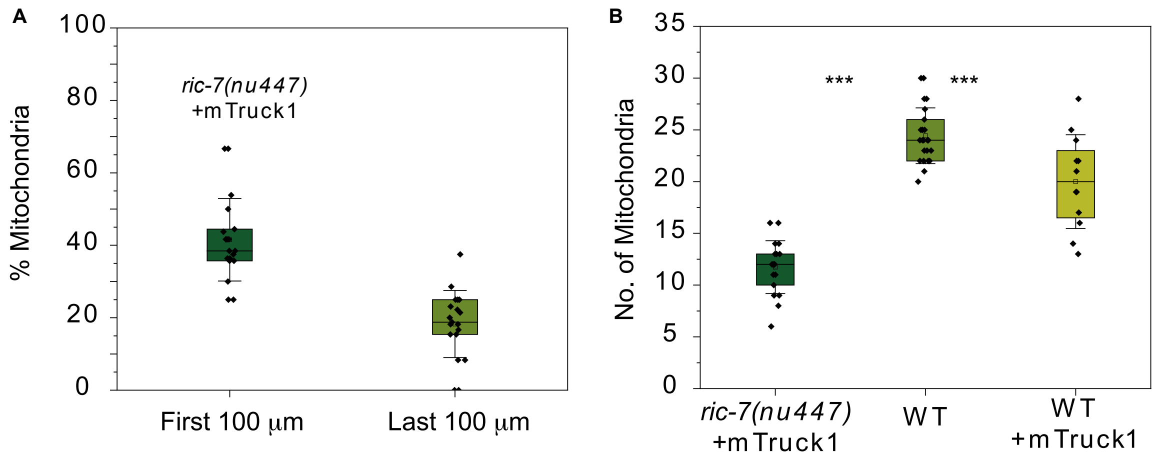


**Supplementary Fig. S10.** ***A***, Percentage of mitochondria present in the first and the last 100 μm length of PLM neuronal process. n >10 animals. ***B***, Number of mitochondria present in PLM neuronal processes after artificial induction of mitochondrial transport in *ric-7(lf)* animals.

**Supplementary Fig. S11.**

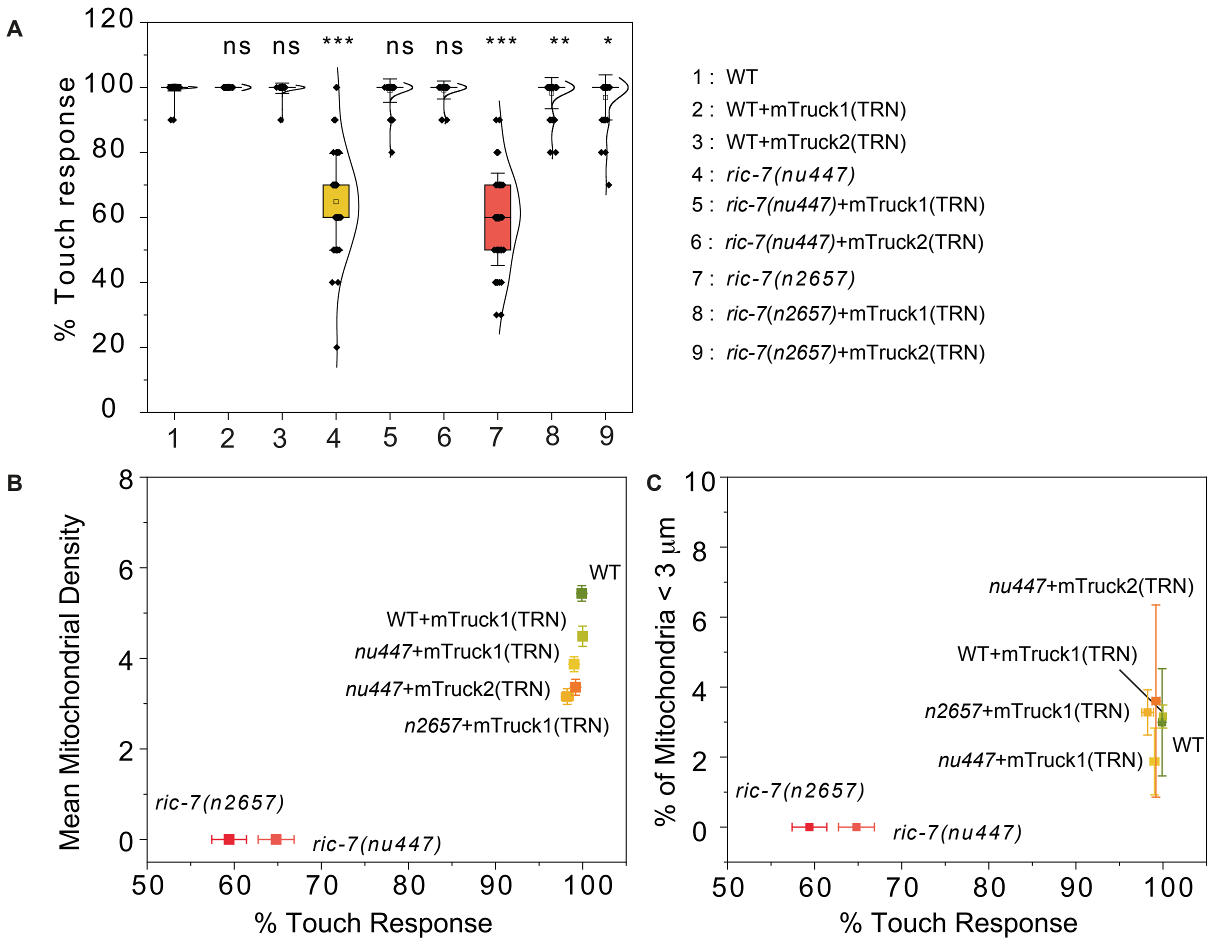


**Supplementary Fig. 11**. ***A***, Violin plot of percentage touch response in WT, *ric-7(lf)*, and *ric-7(lf)* mutant animals expressing mitoTruck. n > 30 animals. ***B-C***, 2D scatter plot of gentle touch response with respect to mean mitochondrial density and percentage of mitochondrial present <3 µm apart in *ric-7(lf)* mutants and in *ric-7(lf)* mutants expressing TRN specific mTruck. For mitochondrial density, percentage of mitochondria present <3µm apart n > 10 animals, for % touch response n > 30 animals.

### Simulation code

```
clear all;
clc;
close('all')
pause(1);
axon=20;

length_max=151.28 + 61.3;
length_prob=61.3*2;

interval=3.0;
fid1 = fopen('axon1.txt','w');
fid2 = fopen('axon2.txt','w');
fid3 = fopen('axon3.txt','w');
figure(1);
x=[];
p=rand(axon,1000);

for j=1:axon
   length(j)=length_max-length_prob*p(j,100);

   N(j)=round(((4.5+1.7)/100)*length(j) - p(j,50)*((1.7*2)/100)*length(j));

   fprintf('axon number-%d  filled with %d mito\n',j,N(j));
   for k=1:N(j)
      x(j,k)=abs(p(j,k))*length(j);
      figure(1);
      plot(x(j,k),j,'r*');
      hold on;
      fprintf(fid1,'%3.0d \t%3.2f\n',j,x(j,k));
   end
   g(j,1:N(j))=sort(x(j,1:N(j)));
   fprintf(fid1,'\n');
   fprintf(fid2,'%3.2f \t %3.2f\t',j,x(j,:));
   fprintf(fid2,'\n');
end
figure(1);
title('Scatter plot of mitochondrial position');
hold off;

%%%% INTER-MITOCHONDRIA DISTRIBUTION  %%%%

for j=1:axon
   for k=1:N(j)-1
      m(j,k)=g(j,k+1)-g(j,k);
      fprintf(fid3,'%3.0d \t%3.2f\n',j,m(j,k));
   end
   fprintf(fid3,'\n');
end

dist_max=length_max/2;
dist_interval=interval;
dist_N=round(0.0 + (dist_max-0.0)/dist_interval);
dist_x=[];
```

```
dist_freq=[];

for z=1:dist_N
   dist_x(z)=(z-1)*dist_interval + dist_interval/2;
   dist_freq(z)=0;
end

for j=1:axon
   for k=1:N(j)-1
      for z=1:dist_N
         if ((dist_x(z)-dist_interval/2) <= m(j,k) && m(j,k) < (dist_x(z)+dist_interval/2))
            dist_freq(z)=dist_freq(z)+1;

         end
      end
   end
end

figure(2);
bar(dist_x,dist_freq);

hold on;
p = (dist_x);
q = (dist_freq);

save('pqfile.txt','p','q');

type('pqfile.txt')
fprintf('pqfile.txt', "%f", p)

plot(dist_x,dist_freq,'r');
xlim([0 100])
set(gca,'xtick',[1 10:10:100])
figure(2);
title('Histogram of the inter-mitochondrial distance');
hold off;

%%%%  MITOCHONDRIA POSITION DISTRIBUTION %%%%

pos_max=length_max;
pos_interval=interval;
pos_N=round(0.0 + (pos_max-0.0)/pos_interval);
pos_x=[];
pos_freq=[];

for z=1:pos_N
   pos_x(z)=(z-1)*pos_interval + pos_interval/2;
   pos_freq(z)=0;
end

for j=1:axon
   for k=1:N(j)
      for z=1:pos_N
         if ((pos_x(z)-pos_interval/2) <= g(j,k) && g(j,k) < (pos_x(z)+pos_interval/2))
            pos_freq(z)=pos_freq(z)+1;
```

```
            end
        end
    end
end

 figure(3);
 bar(pos_x,pos_freq);
 hold on;
 plot(pos_x,pos_freq,'r');
 figure(3);
 title('Histogram of the mitochondrial position');
 hold off;

fclose('all');
```
